# Recent origin of modern clades of iron oxidizers and low clade fidelity of iron metabolisms

**DOI:** 10.1101/2024.09.18.613727

**Authors:** Erik Tamre, Gregory Fournier

## Abstract

Reduced iron was abundant in Earth’s surface environments before their oxygenation, so iron oxidation could have been a common metabolism on the early Earth. Consequently, modern microbial iron oxidation is sometimes seen as a holdover from an earlier biosphere, but the continuity of involved lineages or the metabolic process itself has not been verified. Modern neutrophilic iron oxidizers use cytochrome-porin Cyc2 as the initial electron acceptor in iron oxidation. With the protein as a proxy for the metabolism, we performed a phylogenetic analysis of Cyc2 to understand the evolutionary history of this microbial iron oxidation pathway.

In addition to known iron oxidizers, we identified Cyc2 orthologs in gammaproteobacterial endosymbionts of lucinid bivalves. These bivalves have a robust fossil record and rely on sea-grass meadows that only appear in the Cretaceous, providing a valuable time calibration in the evolutionary history of Cyc2. Our molecular clock analysis shows that extant sampled Cyc2 diversity has surprisingly recent common ancestry, and iron oxidation metabolisms in Gallionellaceae, Zetaproteobacteria, and photoferrotrophic Chlorobi likely originated in the Neoproterozoic or the Phanerozoic via multiple transfer events.

The groups responsible for microbial iron oxidation have thus changed over Earth history, possibly reflecting the instability of niches with sufficient reduced iron. We note that frequent transfer and changing taxonomic distribution may be a general pattern for traits which are selected for sporadically across space and time. Based on iron metabolism and other processes, we explore this concept of a trait’s “clade fidelity” (or lack thereof) and establish its evolutionary importance.

**Importance:** Bacteria can oxidize iron to produce energy. As there was plenty of reduced iron available on the early Earth and there is only a little today, it was sometimes thought that bacteria who oxidize iron today are a small remnant of a larger group that used to do it. We studied the evolutionary history of the iron oxidation pathway that modern bacteria use, and we found that they developed that pathway relatively recently: whoever did it in the past is no longer around today. It would probably be hard for any group of organisms to keep doing iron oxidation over billions of years, since iron availability is so variable: they are likely to go extinct or lose this ability at some point. We suggest this as general trend in evolution that traits which are only sporadically useful are commonly lost – and then re-invented or re-distributed, or the trait will go extinct.

## Introduction

Microbial iron oxidation has been proposed as a common metabolism on the early Earth, given the availability of reduced iron in surface environments before their widespread oxygenation [1]. Banded iron formations (BIFs) present a particularly voluminous body of geological evidence for the abundance of reduced iron once dissolved in the ocean, and photoferrotrophy – one form of microbial iron oxidation – is often implicated in the origin of these formations, given the lack of oxidative power available to oxidize large quantities of iron chemotrophically (e.g., [2], [3]). As BIFs are a quantitatively important presence in the geological record until the Paleoproterozoic, involvement of photoferrotrophy in their creation would suggest a dominant role for that metabolism in early biological productivity. Consequently, modern microbial iron oxidation is sometimes seen as a holdover from a much earlier biosphere [4], yet any continuity of involved lineages or even the metabolic process itself has not been verified. Here, we study the phylogenetic history of Cyc2, a key enzyme in modern microbial iron oxidation, in order to understand the history and antiquity of this pathway as present in modern organisms.

Cytochrome-porin Cyc2 is used by chemolithotrophic Gallionellaceae and Zetaproteobacteria as well as photoferrotrophic Chlorobi as the initial electron acceptor in their iron oxidation pathway [5]: it takes electrons from dissolved iron(II) outside the cell and passes them down an electron transport chain which eventually reaches the terminal oxidase. Since known ecologically important neutrophilic iron oxidizers have Cyc2 and it is highly expressed in environments with a high degree of microbial iron oxidation, it has been proposed as a reliable genomic marker for this metabolism [5]. This allows us to study the phylogenetic history of this protein as a proxy for the history of microbial iron oxidation as observed in modern neutrophilic taxa.

Some previous studies have already considered the possible role of photoferrotrophic Chlorobi in the context of the inference that photoferrotrophy was an important metabolism in the Earth’s past. For example, Thompson et al. [2] propose that they were important players in biogeochemical cycles in the Precambrian, but without explicitly dating the clade or their iron-oxidizing metabolism. Other efforts have estimated the age of photoferrotrophic Chlorobi via molecular dating, with varying results: Ward and Shih [6] found that this clade postdates 300 Ma, while Magnabosco et al. [7] ran analyses with multiple calibration sets which all suggest an age of around 1 Ga. Now that Cyc2 has been identified as the initial electron acceptor in photoferrotrophic Chlorobi as well as other, chemotrophic iron oxidizers, we attempt to date the Cyc2 phylogeny directly as a proxy for the age of the underlying microbial iron oxidation process. Therefore, this phylogenetic history will include the photoferrotrophic Chlorobi and offer an estimate for the age of photoferrotrophy in that clade – which is not necessarily the same as the age of that clade, as the metabolism may not always have been inherited vertically. However, our approach also extends the history of Cyc2-dependent microbial iron oxidation backwards to the last common ancestor of modern Cyc2 sequences – potentially further into the past than the age of any extant group performing iron oxidation today.

## Methods

### Sequence collection and alignment

The Cyc2 sequence in *Chlorobium phaeoferrooxidans* str. KB01 (protein accession number WP_076792910.1) as identified by [8] was used to query the NCBI non-redundant protein database [9] for homologous Cyc2 sequences using BLASTp [10].

Two different datasets of Cyc2 orthologs were assembled, differing in the breadth of the sampling strategy. A broad sample includes the first 500 BLAST hits returned by the search, exhaustively covering the sequence space in and around the main clades of neutrophilic iron oxidizers as well as including any clades containing possible calibration points in the broader protein family. This large sequence set showed multiple suspected misalignments on visual inspection, especially within very distantly related sequences (alignment files in Supplemental Data linked at the end of the manuscript), calling for the cross-validation of results against a smaller dataset including only the descendants of the last common ancestor of Cyc2 sequences in the clades of interest. That narrow sample included a group of 109 sequences from the broad sample (broadly equivalent to what prior work in [5] has called Cluster 1 and already associated with neutrophilic iron oxidation), and a visual inspection of the resulting alignment showed no obvious misalignments. The alignments were made using MAFFT 7.245 [11] with the automatic choice of alignment algorithm (“mafft --auto”).

### Phylogenetic tree search

Based on the alignments, maximum-likelihood phylogenetic trees were built using IQTree 1.6.3 [12]. The ModelFinder function was used to select the best-fitting model, and LG+R7+F (LG substitution model with seven free rate categories and equilibrium frequencies estimated from the data) was selected based on the Bayesian Information Criterion score. The support for each bipartition was estimated using ultrafast bootstraps [13] with 1000 replicates. The relationships between taxa involved in the narrow sampling largely reflected their relationships in the tree based on the broad sampling. Nevertheless, given the potential for misalignments in the broad sample to drive spurious substitution rate inferences, only the best tree obtained for the narrow sampling scheme was used as a basis for molecular clock runs. The tree based on broad sampling was still used as a means of outgroup rooting for the narrow sample.

### Bayesian molecular clock inferences

PhyloBayes 4.1 [14] [15] [16] was used for the relaxed molecular clock runs. Each clock includes a uniform root prior constrained by 2.4 Ga as an older bound, reflective of the dominating presence of aerobic groups on both sides of the root – for example, *Gallionella* and Zetaproteobacteria use oxygen as the electron acceptor for the very iron oxidation process mediated by Cyc2 ( [17] and [18], respectively). As a younger bound, we used 75 Ma, reflecting that all known lucinid bivalves host endosymbionts as adults [19] and the symbiotic relationship with the chemoautotrophic bacteria would have therefore have been established by the lucinid radiation in the late Cretaceous. Thus, as lucinid endosymbiont sequences of Cyc2 represent a subtree of the Cyc2 phylogeny to be dated, the root necessarily predates their divergence. The extreme breadth of the root prior is intended to reflect that lack of prior information on the age of Cyc2-based iron oxidation: that is the question the clock is intended to answer. Nevertheless, note that we reject *a priori* an Archean root, given the topology of the phylogenetic tree, as discussed in the Results section.

It is also important to note that the exact younger bound on the root in the late Phanerozoic does not have a significant impact on the outcome, as the uncertainty in the younger bound selection represents only a small fraction of the total interval covered by the intentionally largely uninformative root prior – in fact, the choice of tree process prior together with the internal node calibration much more dramatically alters the joint root prior than would any small change in the younger bound on the root prior itself (see Supplemental Figure 1). A more extensive discussion of the choice of internal calibrations is given in the Results.

Both the birth-death prior process and a uniform prior on node ages were tested as tree priors, and each relaxed clock model was run with each of the prior processes in PhyloBayes 4.1. We included three clock models: 1) an uncorrelated model with rates on each branch drawn from a gamma distribution (“-ugam”), 2) an autocorrelated model with rates drawn from a log-normal distribution (“-ln”), and 3) an autocorrelated model with rates based on the CIR process (“-cir”). Thus, six clocks in total were run under the posterior, and at least two chains were required to converge for each clock before the posterior distribution would be sampled. In accordance with recommendations in the PhyloBayes manual, convergence was assessed by requiring TRACECOMP values for all estimated variables to be <0.3 and the minimum effective size of the sample to be >50.

Each clock was also run under the prior without the inclusion of sequence information, in order to check the parameters of the joint prior: for example, the impact of the internal node calibrations and the tree process prior means that the joint prior on the root age is considerably narrower than the root prior alone. That also allowed us to distinguish the impact of sequence data from the constraints imposed by the prior. The prior-only runs did not differ between clocks using the same tree process prior, but a different clock model: this is expected, since the only function of the clock model is to convert sequence substitutions into time and sequence data does not impact runs under the prior.

Custom Python scripts were used to read the .datedist files from PhyloBayes, and the ggplot2 package in R was used to visualize the full variability of divergence date estimates in the sample.

## Results

### Phylogenetic tree of Cyc2

The maximum-likelihood tree of Cyc2 protein sequences (Figure 1) does not closely resemble the species tree of the organisms with Cyc2, other than in shallow clades that reflect short periods of largely vertical inheritance close to the present. We observe shallow clades representing subsets of Chlorobi, Betaproteobacteria (in particular, Gallionellaceae), Zetaproteobacteria, Epsilonproteobacteria, Deltaproteobacteria, and Gammaproteobacteria, in addition to scattered sequences of Muproteobacteria, Lambdaproteobacteria, and Nitrospirae.

**Figure 1.**
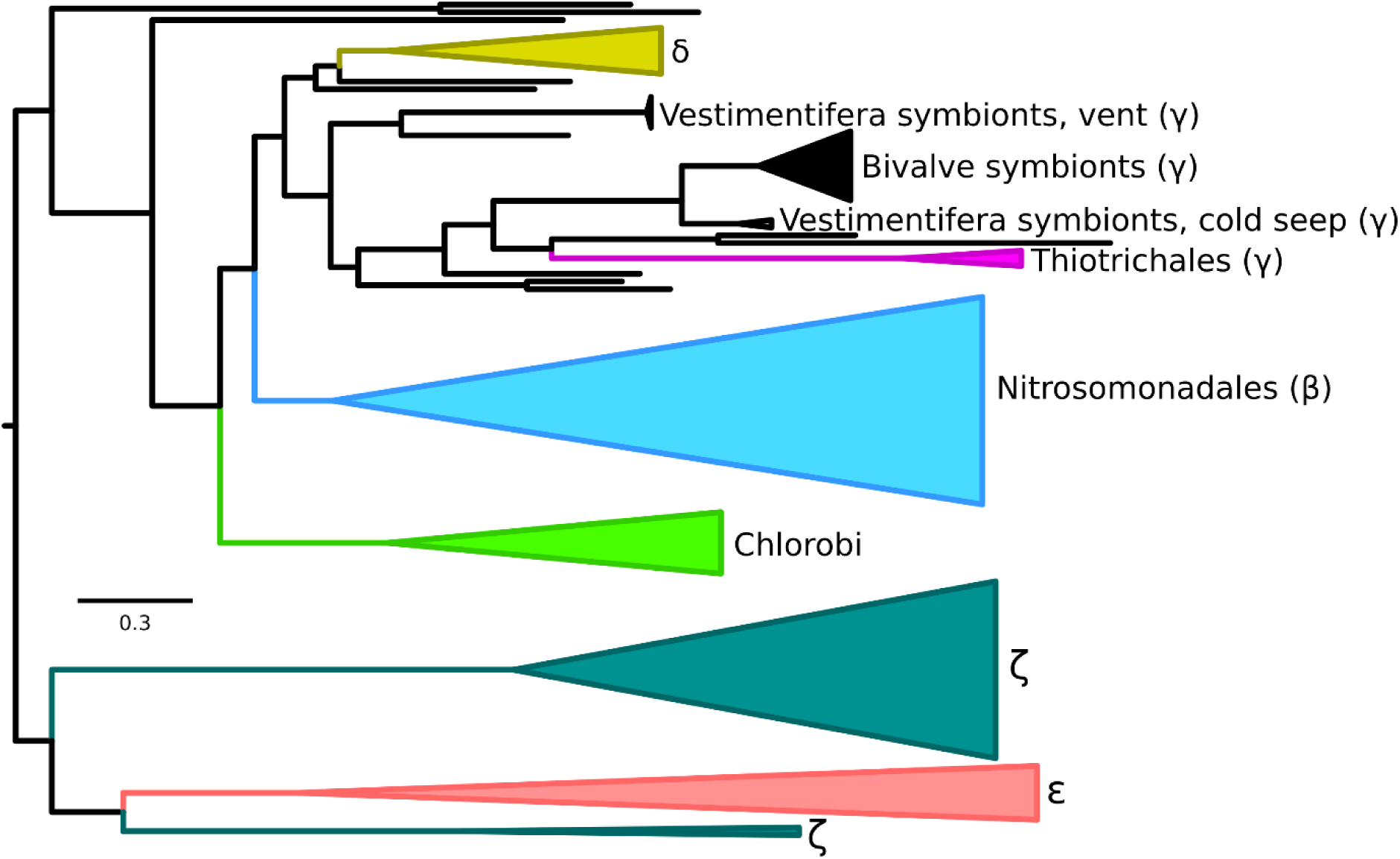
Maximum-likelihood tree of Cyc2 proteins, with groups of sequences in closely related organisms collapsed for clarity. Greek letters refer to taxa falling in the following groups: β – Betaproteobacteria, γ – Gammaproteobacteria, ζ – Zetaproteobacteria (all three now classes in phylum Pseudomonadota); δ – Deltaproteobacteria (now phylum Myxococcota); ε – Epsilonproteobacteria (now phylum Campylobacterota). See Supplemental Figure 2 for the full tree (no groups collapsed), and see Supplemental Data for the annotated tree file.

Even where there are shallow clades where the protein orthologs cluster according to the broad affiliation of the organisms, the relationships between sequences within the cluster often do not reflect the relationships of organisms on the species tree. For example, the gammaproteobacterial sequences fall sister to deltaproteobacterial sequences with betaproteobacterial sequences as an outgroup, whereas Gammaproteobacteria themselves are much more closely related to Betaproteobacteria than either is to Deltaproteobacteria (to the point where the Beta-, Gamma-, and Alphaproteobacteria have recently been renamed as phylum Pseudomonadota, with Deltaproteobacteria classified into their own phylum Myxococcota^1^). Chlorobi are not closely related to any of the Proteobacteria that their Cyc2 sequences group with. Epsilonproteobacteria (recently renamed Campylobacterota) sequences are nested within the zetaproteobacterial sequence diversity, suggesting that the common ancestor of epsilonproteobacterial Cyc2 was transferred into Epsilonproteobacteria from Zetaproteobacteria.

Even within groups of sequences representing one clade of organisms, signatures of horizontal gene transfer are readily observable. Previous work on photoferrotrophic Chlorobi has shown that the tree of Cyc2 sequences in this group does not reflect species tree relationships (compare Fig. 1 and Fig. 2 in [8]); our study reproduces these relationships between Cyc2 orthologs with high support values, which suggest that they are not an artifact of phylogenetic uncertainty due to the relatively small amount of phylogenetic data (in Cyc2, a single protein on average about 400 amino acids in length). Therefore, photoferrotrophs are polyphyletic within Chlorobi not just due to differential loss, but there is also a horizontal gene transfer component.

**Figure 2.**
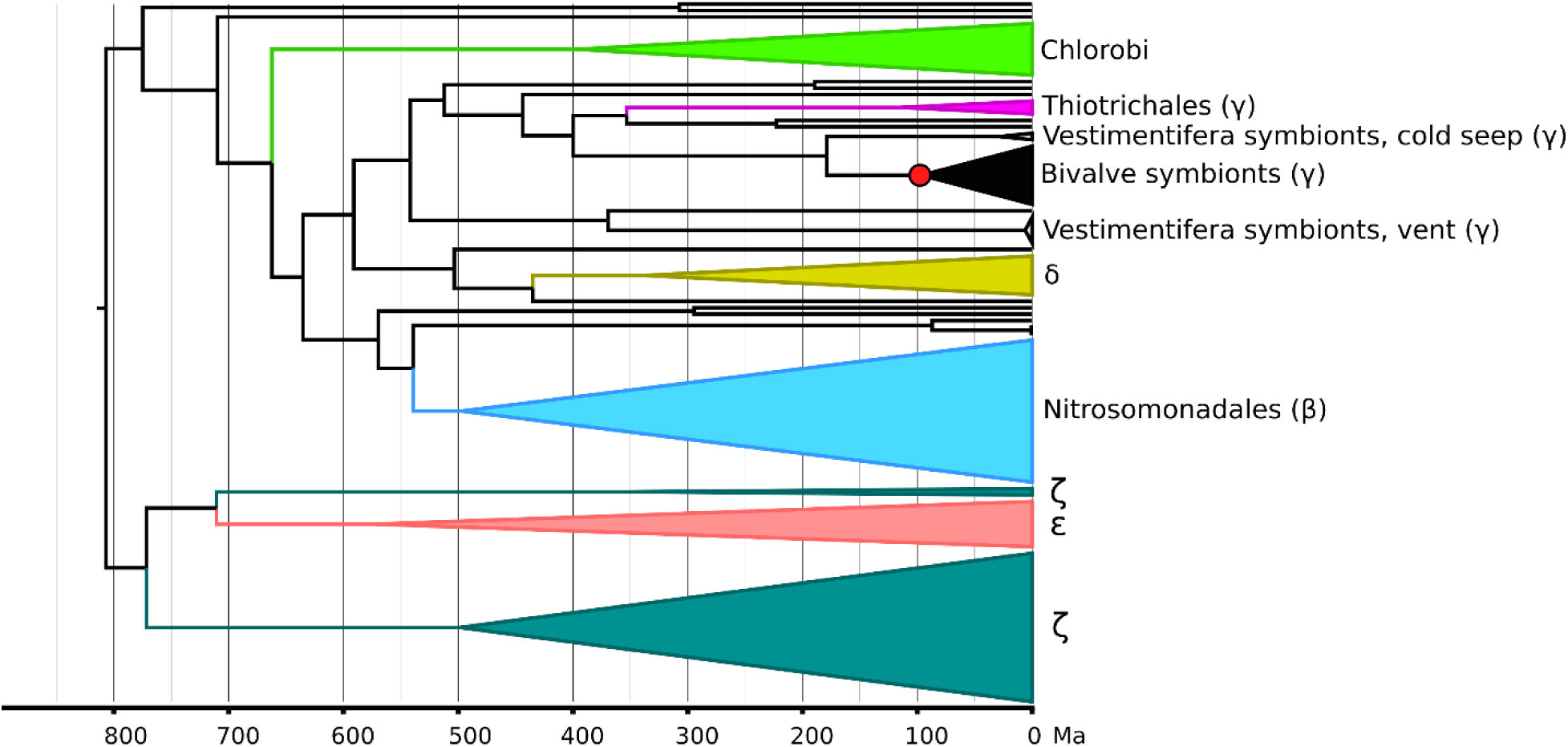
Bayesian relaxed molecular clock based on Cyc2 protein sequences. This representative instance shows mean ages under a uniform tree process prior and an uncorrelated clock model with rates on each branch drawn from a gamma distribution. The red circle marks the calibrated internal node. Greek letters refer to taxa falling in the following groups: β – Betaproteobacteria, γ – Gammaproteobacteria, ζ – Zetaproteobacteria (all three now classes in phylum Pseudomonadota); δ – Deltaproteobacteria (now phylum Myxococcota); ε – Epsilonproteobacteria (now phylum Campylobacterota). See Supplemental Figure 3 for the full tree (no groups collapsed), and see Supplemental Data for the annotated tree file.

The general pattern of non-vertical relationships in the Cyc2 phylogeny suggests a high contribution of horizontal transfer to the overall tree structure. It is also worth noting that the sequences of Cyc2 from gammaproteobacterial endosymbionts of cold-seep vestimentiferan tubeworms are more closely related to sequences from lucinid bivalve symbionts inhabiting a similar environment than either group is to sequences from hydrothermal vent tubeworm endosymbionts. Thus, one possible interpretation of the tree is that we are seeing a history of horizontal gene transfers between organisms inhabiting similar reduced aquatic environments, in addition to periods of vertical inheritance.

### Considerations for dating the tree

When it comes to the interval of time represented by the Cyc2 tree, the first-order observation that groups on the tree represent only shallow clade structure suggests that the interval of time covered by them is relatively short in geological terms. Sequences belonging to major groups cover only a small subset of their diversity, usually dominated by one or two smaller subclades (e.g., *Chlorobium* within Chlorobi, Gallionellaceae within Nitrosomonadales) where lower-level taxonomic assignment is available. While dating these shallow clades is difficult in the absence of a diagnostic paleontological or geochemical record, they are clearly younger than the major bacterial groups they belong to. As they still cover considerable tree depth (about half between the tips and the root for the group sequences belonging to Gallionellaceae, for example), it similarly suggests a relatively young age for the common ancestor of extant Cyc2 diversity.

Similarly, the tree structure alone clearly speaks to the suggestion of any direct relationship between modern photoferrotrophy in phylum Chlorobi and photoferrotrophs in their Archean golden age. Imagine what the tree would look like if the diversity of Cyc2 in extant photoferrotrophic Chlorobi extended all the way to the Archean. In that extreme case, the whole diversity of microaerophilic iron oxidizers such as Gallionellaceae and Zetaproteobacteria should nest within photoferrotrophic Chlorobi on the Cyc2 tree, as their aerobic metabolisms would only be expected to arise starting in the Proterozoic after increased oxygenation of the atmosphere. That is plainly not the tree structure we observe, with the Chlorobi representing only a shallow group on the tree and with all other diversity falling outside of it. This is not a feature of misrooting: see Methods for how the root has been determined, but note also that the shallowness of the group alone makes it very unlikely that the tree should be rooted within this group.

A relaxed Bayesian molecular clock analysis allows for more quantitative testing of these heuristic arguments. Estimating divergence times in the tree requires meaningful prior choices in addition to the sequence data, and these prior choices can be difficult to make in microbial trees with rampant horizontal transfer: tree priors modelling divergence events as consecutive speciation events (such as the birth-death prior) are not necessarily appropriate since many branches traverse multiple lineages through transfer, and calibrations on internal nodes are hard to come by due to a lack of diagnostic fossil or geochemical record in most groups of microbes. Choices of tree prior (the prior on divergence dates of internal nodes) and root prior (the prior on the age of the root) have been discussed above (see Methods), but a more detailed overview of the one applied internal node calibration is appropriate here, since it is itself part of the phylogenetic results.

In addition to iron oxidizers identified in previous studies, we found orthologs of Cyc2 in gammaproteobacterial endosymbionts of lucinid bivalves (Lucinidae) whose radiation is constrained in time by the availability of suitable sea-grass meadows starting in the Cretaceous [20]. Thus, we were able to use the well-dated record of lucinid bivalves [21] to time-calibrate the relevant node in Cyc2 phylogeny. Cyc2 sequences in lucinid endosymbionts group together in the tree, and previous studies have suggested that the corresponding bacterial group forms a monophyletic clade (Fig. 2 in [22]). The most complete phylogenetic and ecological study of this group published – [23], proposing the group as “Ca. Thiodiazotropha” – only observed its members in symbioses with lucinid bivalves. While endosymbionts are acquired from the environment in lucinid bivalves and there is no faithful vertical co-evolution between the host and the symbiont at the species level [19], this evidence suggests that members of the group specifically enter into symbioses with lucinid bivalves and thus that their last common ancestor was also a lucinid endosymbiont. That allows us to impose a lower bound on the node representing this last common ancestor: it should not pre-date the appearance of lucinid bivalves. In a previously published molecular clock of Lucinidae well constrained by fossil calibrations, the divergence of Lucinidae from its sister group takes place 170 million years ago [21]. Given that the establishment of the symbiotic relationship could in principle have taken place in stem lucinids, we use this total group age for Lucinidae as a conservative older bound for the age of the last common ancestor of lucinid endosymbiont Cyc2 sequences.

### Molecular clock outcomes and sensitivity testing

In runs under the posterior (see Table 1 for a summary of key node ages and Figure 2 for a representative clock), the most general pattern is that clocks using a birth-death prior rather than a uniform prior on divergence dates produce significantly older ages for internal nodes. This is likely because the node constrained by the internal calibration appears much shallower in the tree (i.e., at lower relative tree depth) under the birth-death prior, and the full depth of the tree is therefore estimated to be greater in absolute terms. For example, the mean age of the last common ancestor of Cyc2 sequences in photoferrotrophic Chlorobi falls between 270 and 400 Ma for runs with the uniform prior, but between 530 Ma and 625 Ma for runs with the birth-death prior. However, for the purposes of testing the hypothesis of early origin for this group, it is crucial to observe that under no model does a significant fraction of the probability mass exceed 1 Ga: see Figure 3 for full posterior probability distributions for the age of this node under all models. In other words, even though the tree prior process considerably impacts ages of internal nodes (likely due to the relative lack of sequence data, as Cyc2 has about 400 amino acids in most organisms), its influence on younger bounds of node ages is more profound than on older bounds.

**Figure 3.**
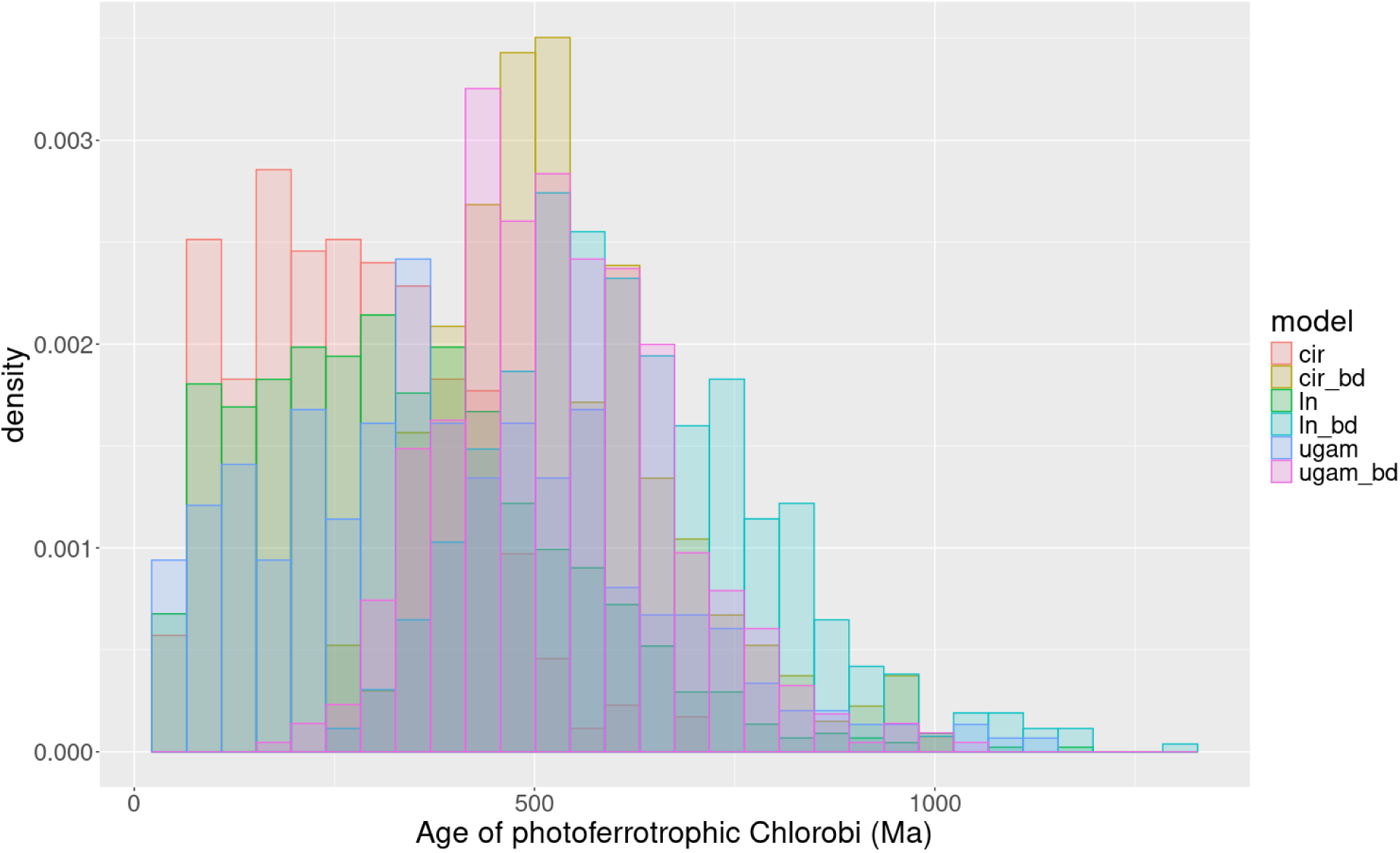
Overlaid posterior probability distributions for the age of the last common ancestor of Cyc2 sequences in photoferrotrophic Chlorobi under all combinations of clock models and tree prior processes, summarizing the full range of dating uncertainty. Clock models: “cir” – autocorrelated clock with branch rates based on the CIR process; “ln” – autocorrelated clock with branch rates drawn from a log-normal distribution; “ugam” – uncorrelated clock with branch rates drawn from a gamma distribution. Clocks with “bd” have been run under a birth-death prior, while those without that marker use a uniform tree process prior.

**Table 1.**
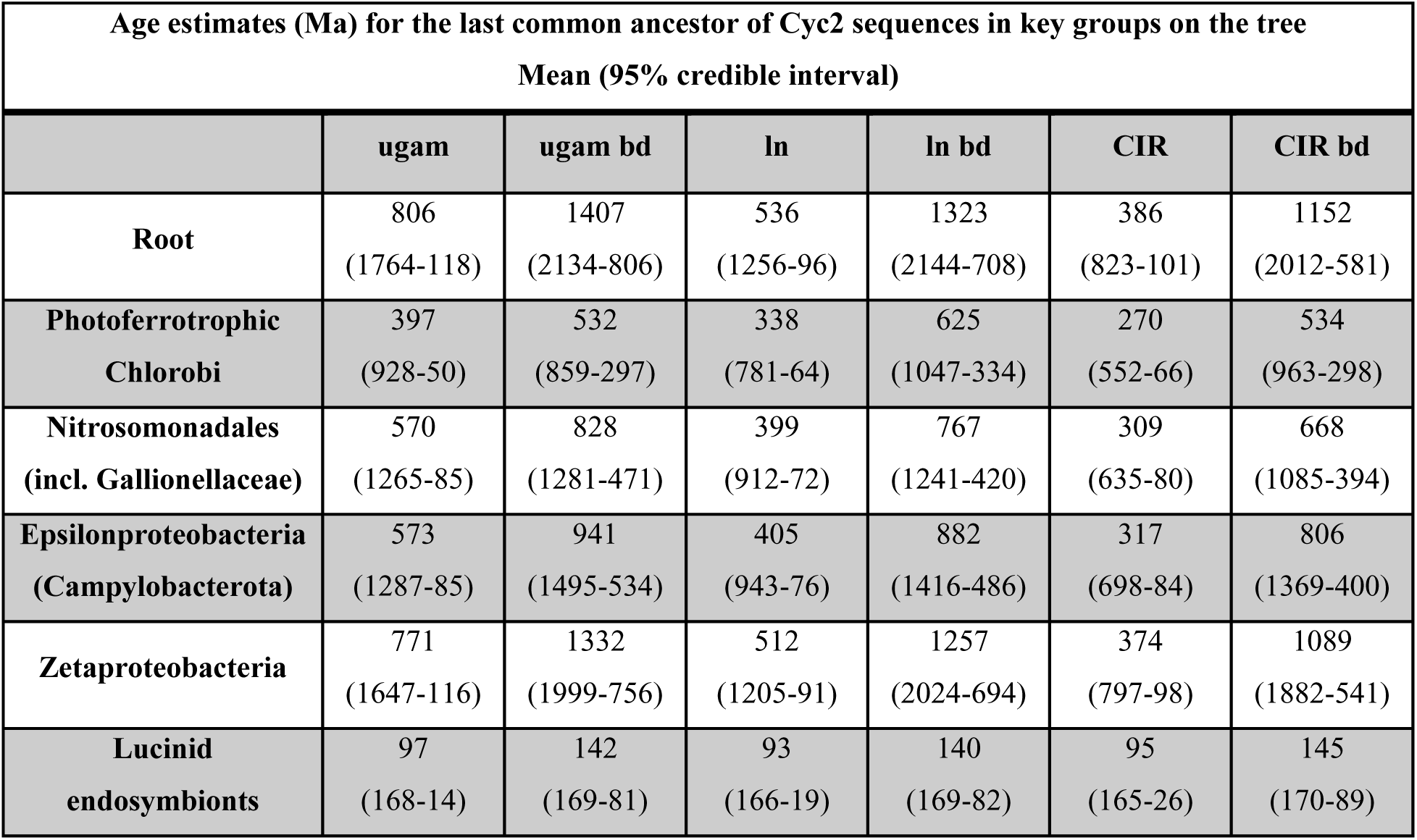
A summary of age estimates for key nodes on the Cyc2 tree. Note that these estimates are not for the last common ancestor of organisms in each group, but rather the last common ancestor of Cyc2 sequences present in members of the group. Clock models: “ugam” denotes an uncorrelated model with rates on each branch drawn from a gamma distribution, “ln” denotes an autocorrelated model with rates drawn from a log-normal distribution, and “cir” denotes an autocorrelated model with rates based on the CIR process. Addition of “bd” signifies that the clock was run under the birth-death tree process prior; otherwise the uniform prior was used.

In general, greater uncertainties on younger bounds than older bounds are to be expected, as the one internal node calibration used only constrains its older bound: the substitution rate is constrained to be above a certain value and does not allow branches to stretch arbitrarily far into the past. This offers a one-sided test for the hypothesis of ancient origin for the iron oxidation process in question. Note also that the presence or absence of correlation between substitution rates on adjacent branches has less of an impact on divergence dates than the prior process. That observation is particularly important since we have imposed a calibration on a clade of endosymbionts, where substitution rates could substantially differ from those in related free-living lineages. Autocorrelated and uncorrelated clocks have a different capacity to take such a transition into account, and it should increase overall confidence in these dating estimates that outcomes of autocorrelated and uncorrelated clocks do not contrast greatly with each other.

Other clades of modern iron oxidizers on the tree show patterns similar to photoferrotrophic Chlorobi. Mean ages of 660-830 Ma are calculated for the last common ancestor of Gallionellaceae sequences under models with a birth-death prior, and 300-570 Ma under models with a uniform prior. For zetaproteobacterial Cyc2 sequences, whose last common ancestor stretches deeper into the tree, the age estimates range between 370-780 Ma for the uniform prior and between 1080-1340 Ma for the birth-death prior. Full chronograms showing the 95% credible intervals for all internal nodes are shown as Supplemental Figures 3-8.

## Discussion

The relaxed molecular clock analysis shows that iron oxidation with Cyc2 in modern photoferrotrophic Chlorobi is a relatively young trait, originating in the late Neoproterozoic or the Phanerozoic. This result also agrees with previous estimates of relatively recent origin for the clade of Chlorobi in which photoferrotrophy is present. It is inconsistent with views of the current process of photoferrotrophy as a direct descendant of photoferrotrophic iron oxidation in the Archean and Paleoproterozoic that may have played a role in the deposition of extensive BIFs during these times.

Furthermore, timing estimates for the last common ancestor of Cyc2 sequences in other major microbial iron oxidizers such as Gallionellaceae and Zetaproteobacteria are also relatively recent across all clock runs, with the earliest likely emergence of Cyc2-based iron oxidation in Neoproterozoic in Gallionellaceae and in Mesoproterozoic in Zetaproterobacteria. Even though fossils morphologically similar to modern chemolithotrophic iron oxidizers date back to at least the Paleoproterozoic (e.g., [24] & [25]), the recent origin of the modern clades suggests that these early fossils do not represent known extant groups. While chemolithotrophic iron oxidation and photoferrotrophy as metabolisms are undoubtedly very ancient, the taxonomy of responsible organisms has changed profoundly through time.

In our effort to date the history of Cyc2, we have chosen to be conservative in the selection of priors and inclusive in testing the impact of a broad variety of molecular clock models on the divergence date estimates. This is to recognize that different clocks often yield very different age estimates, and we do not have a good way to argue for the use of one set of parameters over another for the very reason why ages in microbial natural history are hard to constrain: rock record calibrations are scant and our understanding of the underlying evolutionary process (impact of transfer vs. vertical inheritance, estimate of substitution rates across branches in the distant past) is limited. In this study, dating outcomes hinge most significantly on the choice of the tree process prior: the birth-death prior models nodes as consecutive speciation events (though branches in this tree commonly traverse distantly related lineages owing to transfer), while the uniform prior assumes no knowledge of evolutionary process whatsoever and is sometimes dismissed because there is no known evolutionary process that would yield such a distribution of branching events [26]. We have no way to decide between them, and thus we have to live with considerable dating uncertainties. Nevertheless, the finding that modern iron oxidation is quite young, also suggested by the shallow clade structure of the tree alone, is robust to extensive sensitivity testing carried out on formal molecular clocks.

Ecologically, the shallow clade depths could reflect the environmental history and bioavailability of iron: in a largely oxygenated world, niches with sufficient reduced iron are geographically restricted and ecologically fragmented, and they do not persist over geologically long periods of time. After a while, this electron donor becomes scarce and organisms either adapt by loss of the pathway, or they simply leave no descendants. However, there are always refugia somewhere in the ocean with sufficient reduced iron, even if an individual instance of that environment does not last particularly long. When reduced iron becomes locally abundant, Cyc2 (and potentially other machinery for iron oxidation) is shared by horizontal transfer by the bacterial populations in this environment, and it is broadly retained due to the useful metabolic trait it imparts. It is possible that retention of iron oxidation machinery would have been more consistent and patterns of vertical inheritance more pronounced if we examined their phylogeny in the Archean, when selection for being able to use reduced iron as an electron donor would have been more consistent in an ocean with plenty of reduced iron.

If the ecology of iron metabolism in the recent oceans is defined by its spotty availability, this could also be reflected in the evolutionary history and phylogenetic distribution of other traits partaking in iron metabolism. While Cyc2-dependent iron oxidation is useful where reduced iron is available, siderophores are commonly used for scavenging iron where it is unavailable [27]. If the presence of this reverse condition is also somewhat variable in space and time, then loss, transfer, and re-acquisition of siderophore biosynthesis pathways could show similar evolutionary patterns. Indeed, for example in Cyanobacteria, siderophore biosynthesis pathways show a patchy distribution: the production of a given siderophore is often polyphyletic, and even closely related taxa can produce different siderophores [28]. Though reconstructing phylogenetic histories for particular siderophores is challenging owing to the promiscuity of the biosynthesis enzymes [28], these observations are definitely consistent with frequent departures from vertical inheritance. Shallow clade depth and frequent radiation, loss, and transfer may characterize a general pattern in the evolution of iron metabolism machinery.

Regarding the ecological dynamics influencing the evolutionary history of iron utilization, it is worth adding that many of the organisms appearing in this study of Cyc2 also seem to oxidize sulfur. For example, the gammaproteobacterial endosymbionts within lucinid bivalves that were essential for dating this history have been previously identified as carrying out sulfur oxidation for their hosts: to our knowledge this study is the first report of the presence of Cyc2 in these endosymbionts. Though this is not an issue for dating purposes as our approach is agnostic of the function of Cyc2 in these microbes, it has not been experimentally demonstrated to carry out iron oxidation in them. Beyond the gammaproteobacterial endosymbionts of lucinid bivalves, the green sulfur bacteria (Chlorobi) on the tree are so named because they most commonly rely on sulfide as an electron donor for photosynthesis, and the Epsilonproteobacteria are largely sulfur-oxidizing chemotrophs. In future work, it may be illuminating to consider further the reasons behind the taxonomic overlap between metabolisms utilizing reduced iron and sulfur: is it simply an ecological reflection of organisms living in environments with reduced fluids where each species is present, or is the reason a more profound overlap between the adaptations and machinery necessary for these metabolisms? These questions will become simpler to answer once we understand more clearly how the evolutionary histories of iron and sulfur metabolisms are separate or intertwined.

In describing the evolutionary and phylogenetic history of these metabolisms, we propose defining a trait’s “clade fidelity” [29]: its tendency towards vertical inheritance and a stable taxonomic distribution over time, as opposed to frequent transfer and change in its taxonomic distribution. Microbial iron oxidation considered here is a clear example of a trait with low clade fidelity: throughout its history, it is frequently transferred between clades or lost and regained. Some clades with the trait likely become extinct while new clades become associated with it, and at any given time the trait’s history in contemporary clades is shallow. For comparison (Figure 4), this is in striking contrast to a trait like oxygenic photosynthesis, which evolved once in an ancestor of modern cyanobacteria and is exclusively present in the descendants of this organism, having been faithfully vertically inherited for well over two billion years [30].

**Figure 4.**
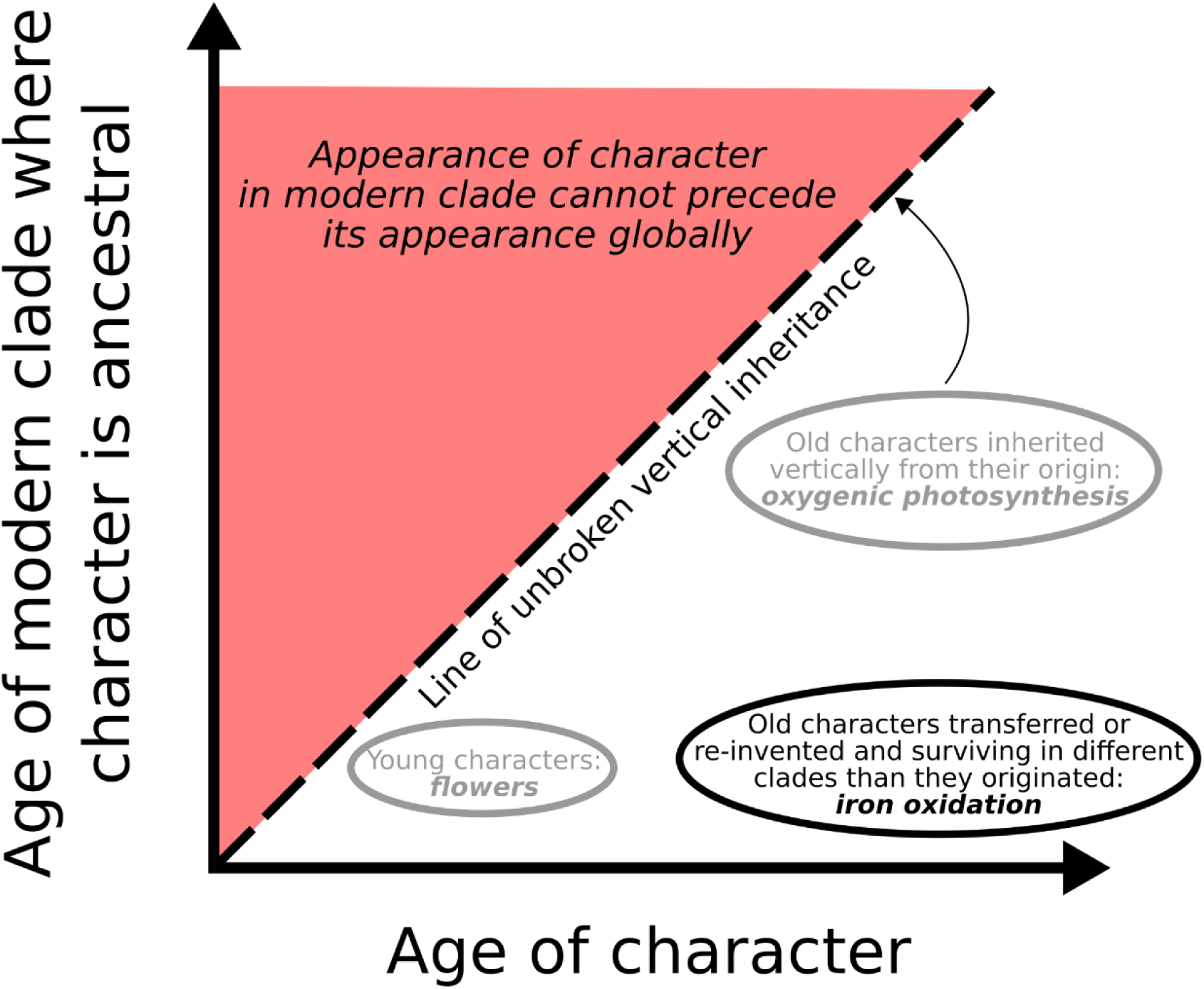
A schematic depicting examples of traits with varying levels of clade fidelity. The clade fidelity of a character is defined by its vertical distance from the 1:1 line (dashes) compared to its distance from the x-axis. Characters with low clade fidelity (such as microbial iron oxidation) plot close to the x-axis, as modern clades with the character are much younger than the character itself; characters with high clade fidelity (such as oxygenic photosynthesis) plot closer to the 1:1 line or even onto it. Note that it makes sense to talk about the clade fidelity of a character mainly over long timescales where radiation, extinction, and transfer is likely to occur. In the microbial world (such as for iron oxidation), geological evidence of a trait’s presence is often necessary to infer its age, as evolutionary inferences made using the genetic record in extant organisms do not give access to extinct lineages.

On a planetary scale, there may be some selection for low clade fidelity on traits whose usefulness is episodic. A type of iron oxidation we considered in particular detail is photoferrotrophy, and it is interesting to note that the finding of its relative youth as carried out in the modern by some green sulfur bacteria parallels the relatively shallow histories of other forms of anoxygenic photosynthesis (for a summary, see [30]). Unlike oxygenic phototrophs, anoxygenic phototrophs are limited by the variable abundance of the electron donor. It is striking, therefore, that forms of anoxygenic photosynthesis would show low clade fidelity, whereas oxygenic photosynthesis with its more reliable substrate exhibits highest possible clade fidelity, having only evolved once and never known to have been transferred to another microbial lineage. This is not to say that the reliable abundance of substrate causes high clade fidelity, but rather that a metabolism with high clade fidelity might not be able to survive unless consistently favored over geological time, including by the abundance of substrate. There may have been many forms of photoferrotrophy in the past that, for whatever reason (complex underlying machinery, lack of modularity or potential to be integrated with other metabolisms), showed high clade fidelity – but these forms would be more likely to go extinct given their limited resilience to contexts where photoferrotrophy was not favored. A form of photoferrotrophy with low clade fidelity, distributed in a set of organisms and localities, is more likely to survive and subsequently radiate again when favorable conditions emerge.

Such a model assumes that selection operates not just on organisms themselves, but also on biotic processes. This view has recently been popularized by the “It’s the song not the singer” theory initially proposed by Ford Doolittle and colleagues in the context of host-symbiont systems [31], but subsequently generalized to broader interactions between organisms in the ecosystem and even biogeochemical cycles [32]. This theory emphasizes that interactions between organisms or processes mediated by them (“songs”) often outlast individual clades (“singers”) and should be considered proper units of selection themselves. If that is the case, it is easy to imagine clade fidelity as one of the main aspects of a trait or process to be under selection. Processes with high clade fidelity are contingent on the survival of individual clades in which they occur and both more likely to suffer from extinction and less likely to re-emerge convergently thereafter.

From a planetary perspective, the clade fidelity of a trait could even have implications for the robustness of the biosphere in which that trait participates. This is easiest to imagine if the trait is a metabolic ability to carry out a biogeochemical process which participates in a web of biotic interactions. It thus influences other organisms as well as the collective they make up together – the biosphere as a whole, whose stability over Earth history may have included a small positive contribution from the low clade fidelity of microbial iron oxidation.

To summarize the history of microbial iron oxidation from the Archean to the present in Doolittle’s terms, the singers have changed, but the song has endured. It is uncertain whether Cyc2 itself played a role in microbial iron oxidation before the emergence of the groups currently using it: Cyc2 may have been carried in the more distant past by lineages that left no extant descendants, yet transferred their iron oxidation machinery into lineages that did. Alternatively, earlier iron oxidation machinery could have been unrelated. Either way, when considering microbial iron oxidation in the geological past, it needs to be assumed that it transcended its modern taxonomic distribution – for traits of low clade fidelity, looking to modern organisms as proxies of past process is particularly perilous.

## Supplemental Data

The Supplemental Data files for this manuscript, including all alignment, tree, and chronogram files, can be accessed at https://doi.org/10.5061/dryad.905qfttvf.

## Supplemental Figures

**Supplemental Figure 1.**
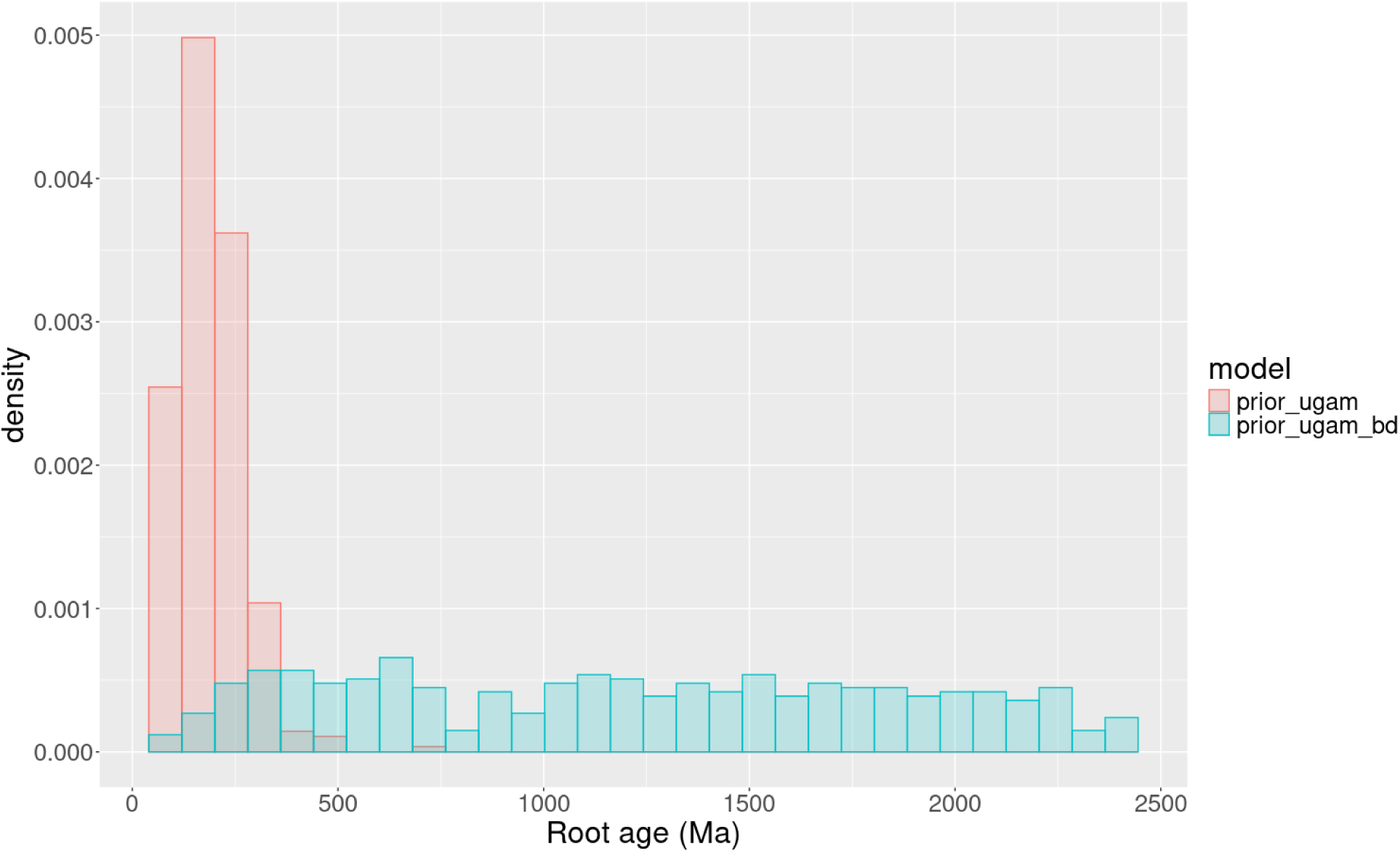
Root priors conditional on the internal node calibration and the tree process prior (uniform vs. birth-death prior on divergence dates).

**Supplemental Figure 2.**
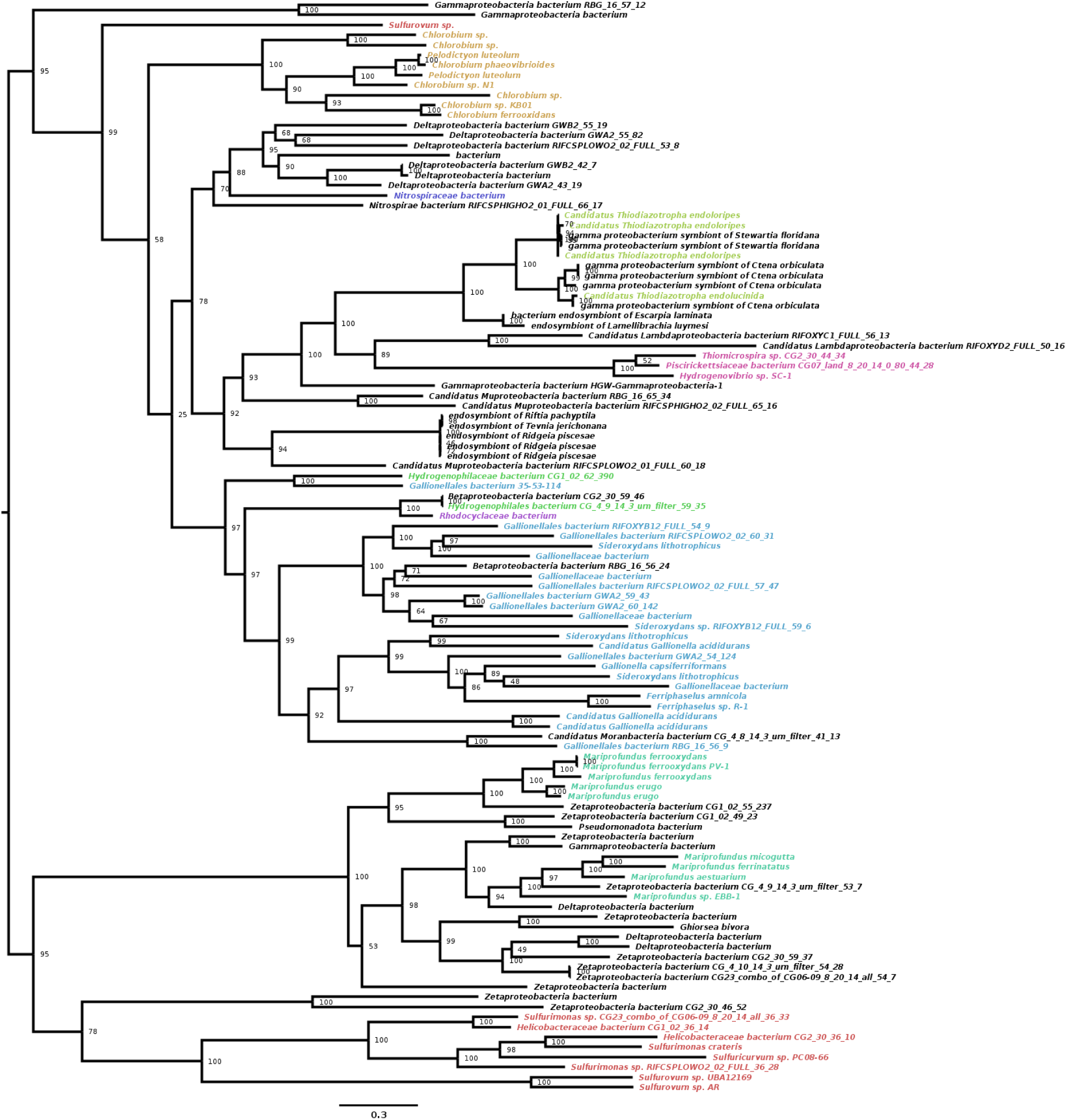
Full maximum-likelihood phylogenetic tree of Cyc2 sequences. Node labels represent ultrafast bootstrap support for the labeled bipartition. Taxa are colored according to taxonomic order (where assigned to an order).

**Supplemental Figure 3.**
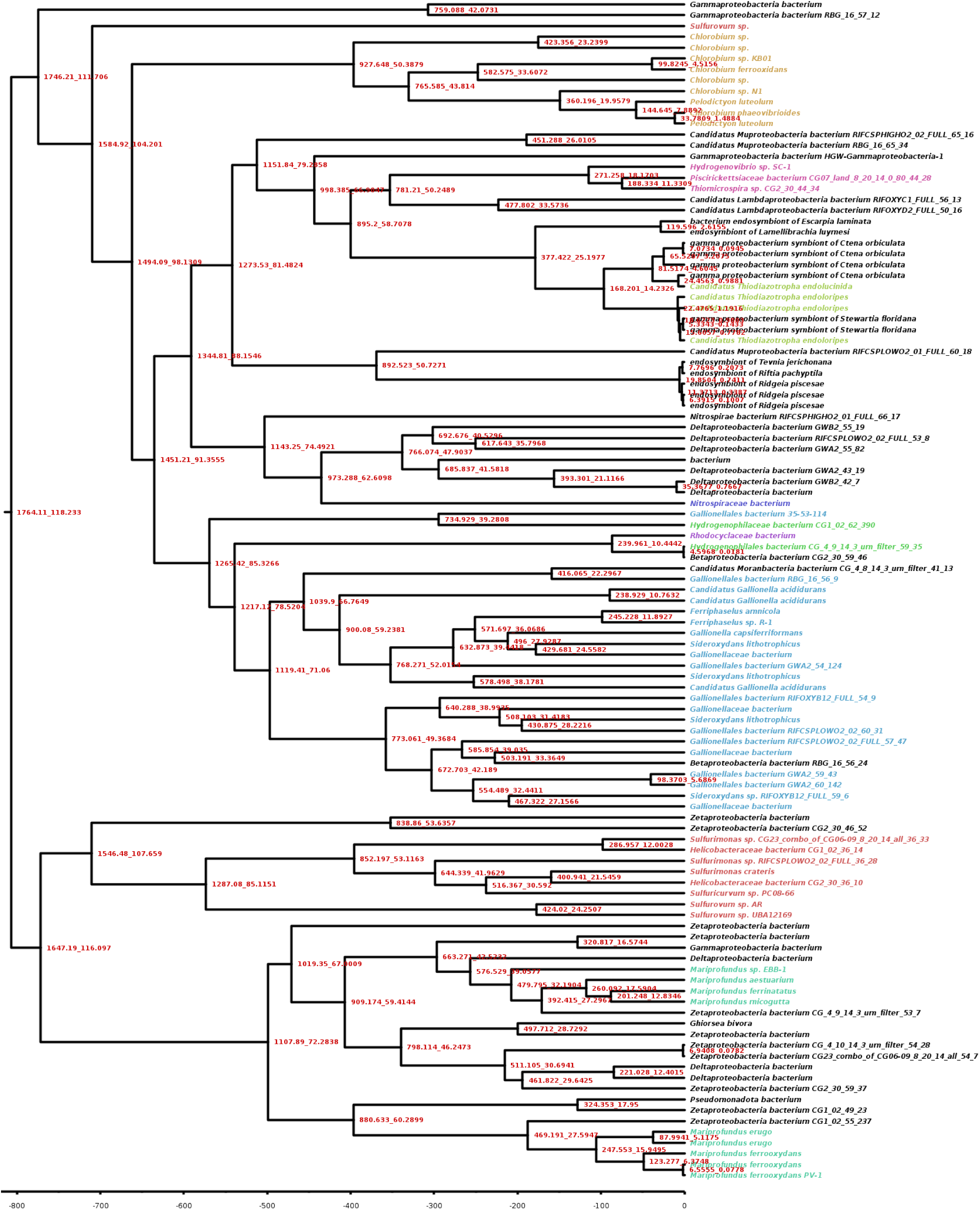
Full chronogram of Cyc2 sequences built using the uniform tree process prior and the uncorrelated gamma-distributed clock model. All ages in Ma. Node labels represent 95% posterior credible intervals for node age. Taxa are colored according to taxonomic order (where assigned to an order).

**Supplemental Figure 4.**
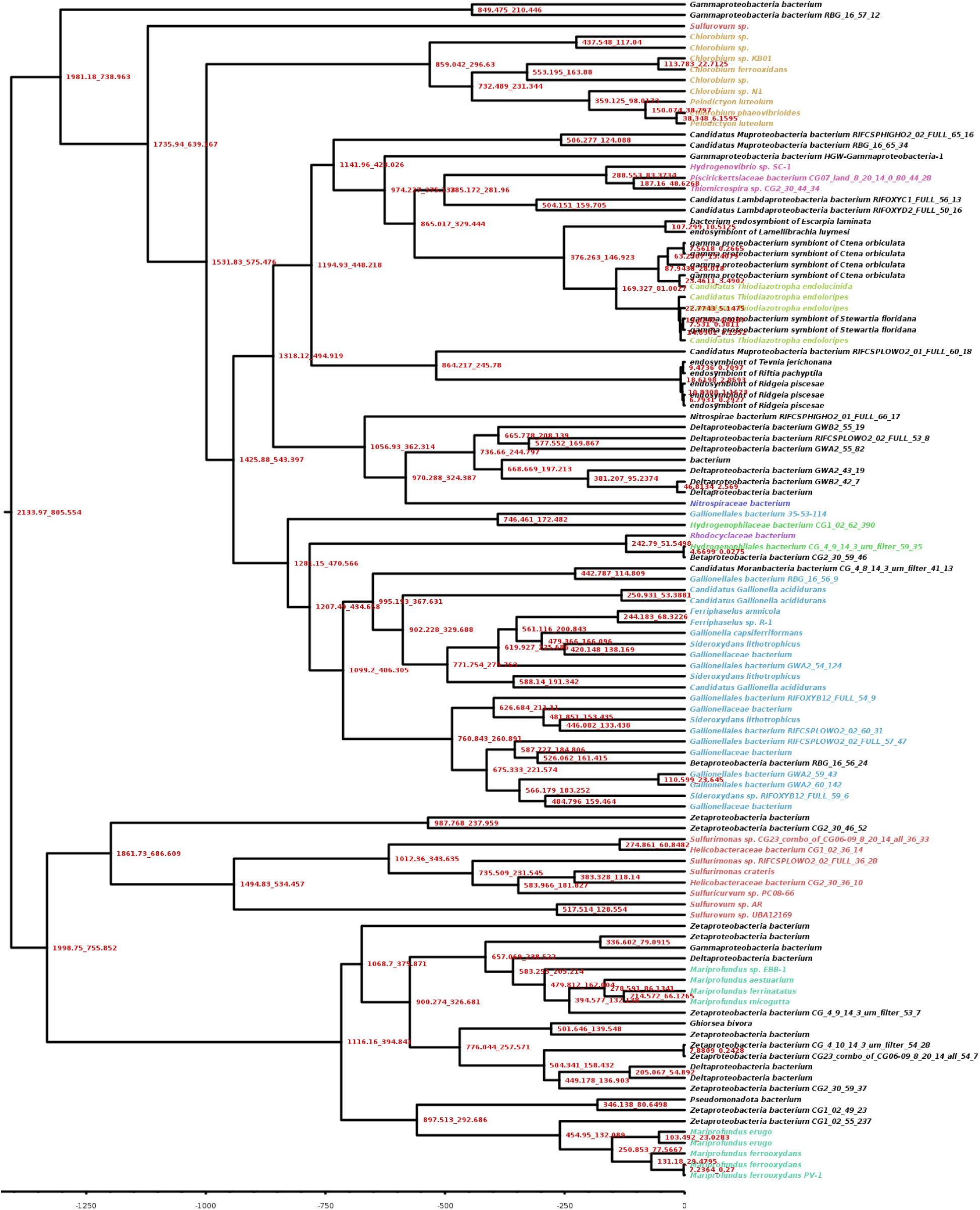
Full chronogram of Cyc2 sequences built using the birth-death tree process prior and the uncorrelated gamma-distributed clock model. All ages in Ma. Node labels represent 95% posterior credible intervals for node age. Taxa are colored according to taxonomic order (where assigned to an order).

**Supplemental Figure 5.**
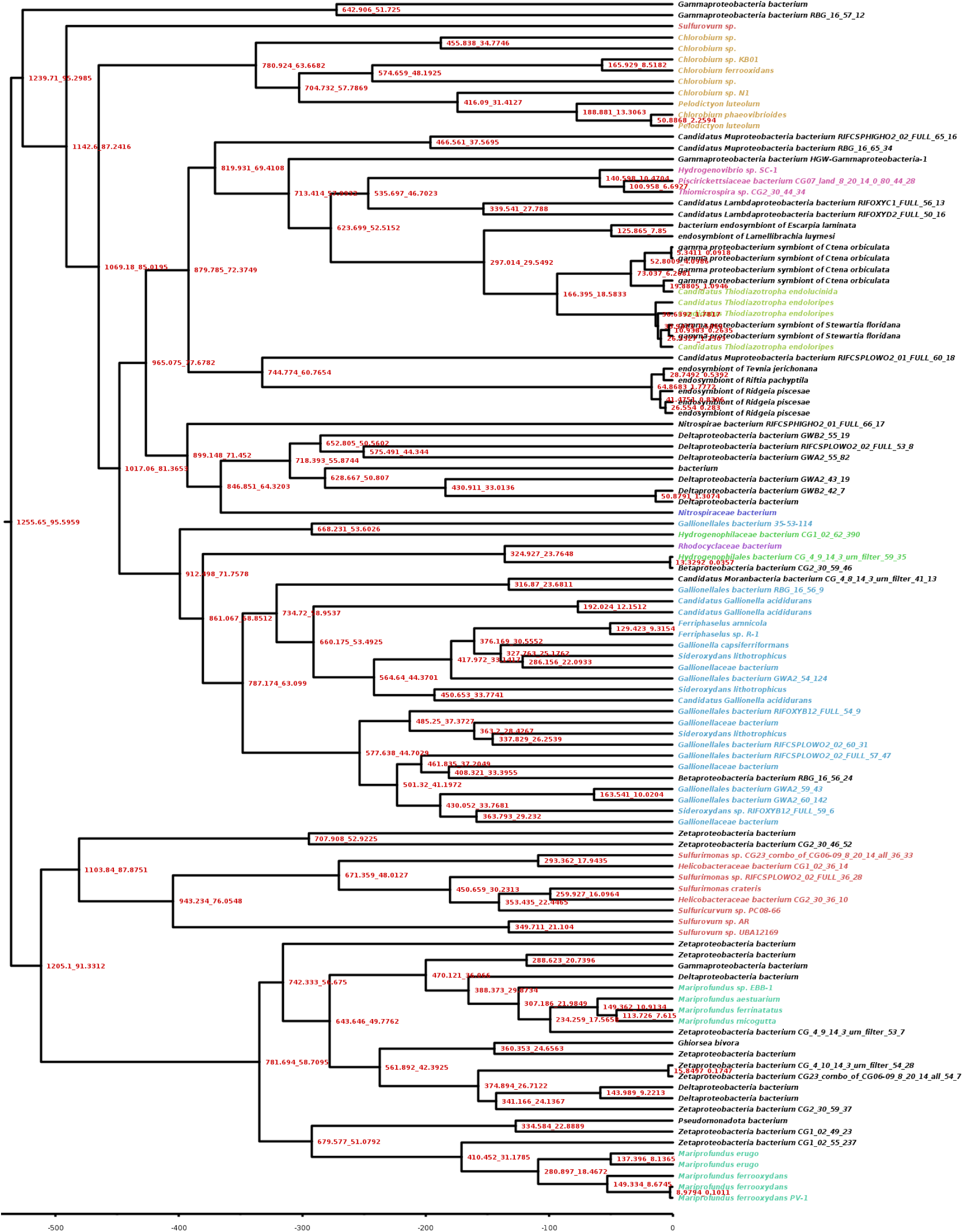
Full chronogram of Cyc2 sequences built using the uniform tree process prior and the log-normal clock model. All ages in Ma. Node labels represent 95% posterior credible intervals for node age. Taxa are colored according to taxonomic order (where assigned to an order).

**Supplemental Figure 6.**
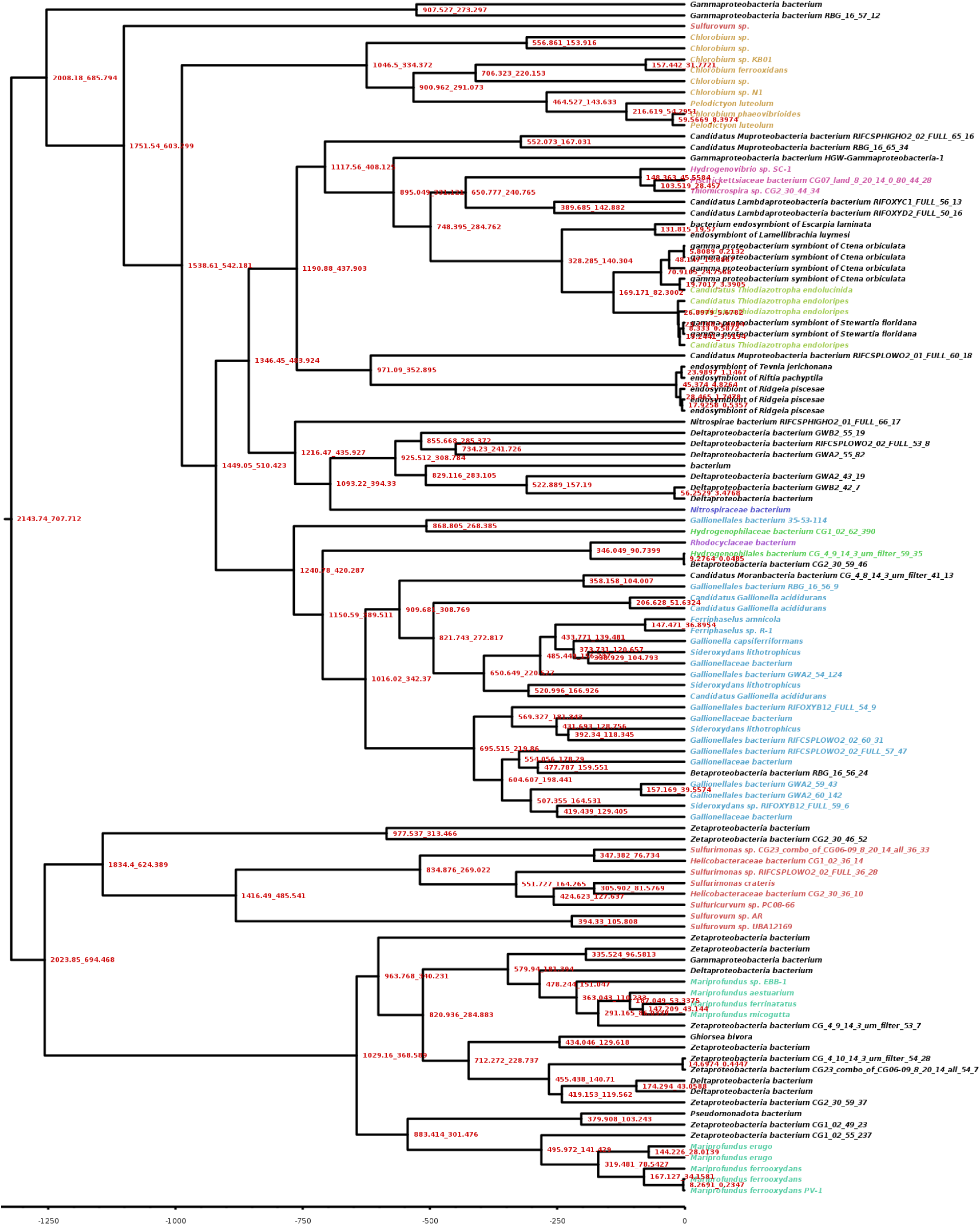
Full chronogram of Cyc2 sequences built using the birth-death treeprocess prior and the log-normal clock model. All ages in Ma. Node labels represent 95% posterior credible intervals for node age. Taxa are colored according to taxonomic order (where assigned to an order).

**Supplemental Figure 7.**
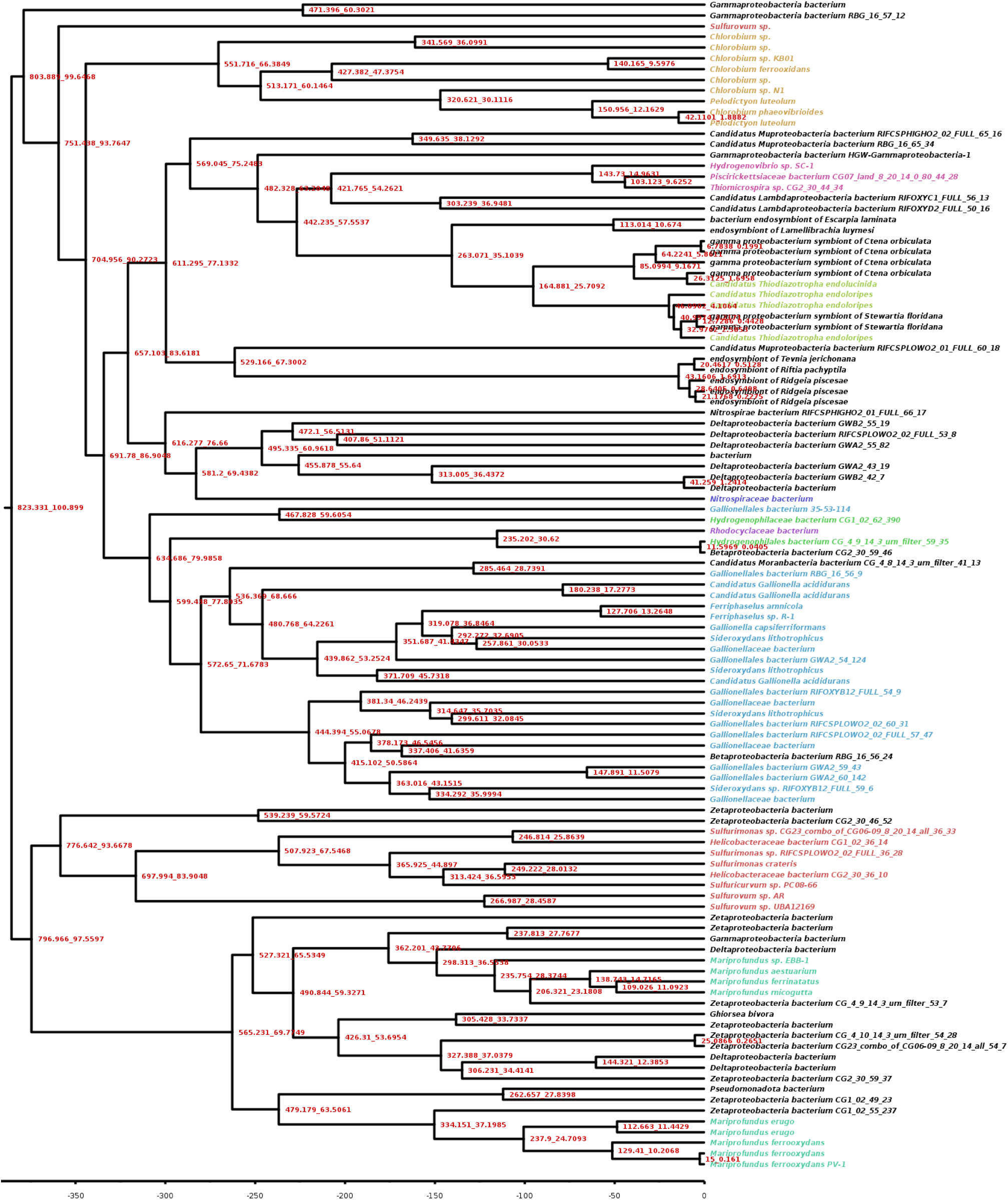
Full chronogram of Cyc2 sequences built using the uniform tree process prior and the CIR clock model. All ages in Ma. Node labels represent 95% posterior credible intervals for node age. Taxa are colored according to taxonomic order (where assigned to an order).

**Supplemental Figure 8.**
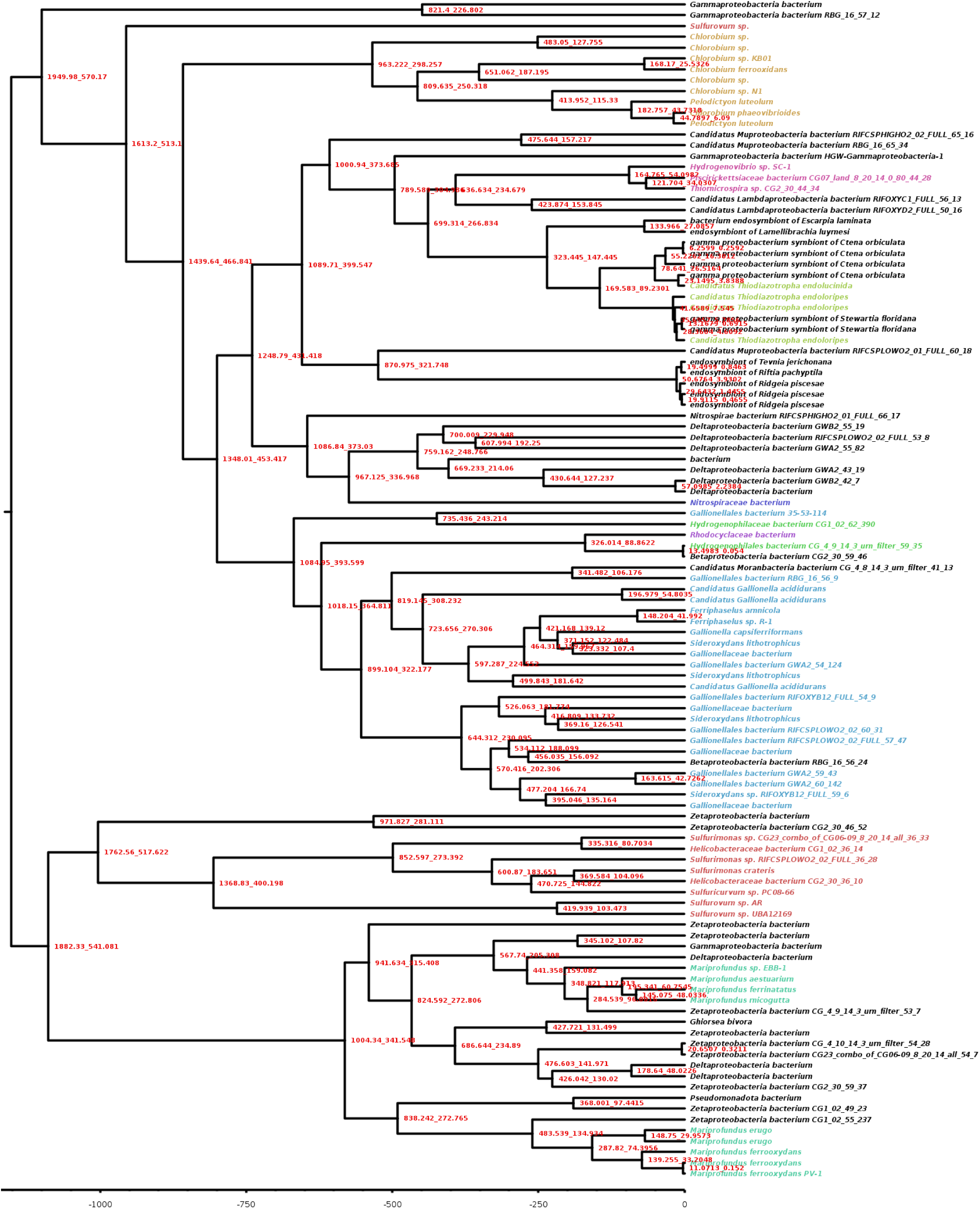
Full chronogram of Cyc2 sequences built using the birth-death tree process prior and the CIR clock model. All ages in Ma. Node labels represent 95% posterior credible intervals for node age. Taxa are colored according to taxonomic order (where assigned to an order).

## Footnotes

1 We refer to bacterial clades using the nomenclature before the changes implemented in 2021 but subject to continuing contention [33] [34]. This is largely motivated by the lingering use of the old nomenclature in the NCBI Taxonomy Database as well as in taxon names for sequences uploaded before the change, which make up the vast majority of the sequences in the trees and need to be reconcilable to the main text. Where applicable, we still mention the recent classification upon the first appearance of the clade in the text to allow for reconciliation with the new nomenclature as well.

